# Single molecule analysis of effects of non-canonical guide RNAs and specificity-enhancing mutations on Cas9-induced DNA unwinding

**DOI:** 10.1101/642223

**Authors:** Ikenna C. Okafor, Digvijay Singh, Yanbo Wang, Minhee Jung, Haobo Wang, John Mallon, Scott Bailey, Jungjoon K. Lee, Taekjip Ha

## Abstract

Cas9 has made a wide range of genome engineering applications possible. However, its specificity continues to be a challenge. Non-canonical gRNAs and new engineered variants of Cas9 have been developed to improve specificity but at the cost of the on-target activity. DNA unwinding is the primary checkpoint before cleavage by Cas9 and was shown to be made more sensitive to sequence mismatches by specificity-enhancing mutations in Cas9. Here we performed single-molecule FRET-based DNA unwinding experiments using various combinations of non-canonical gRNAs and different Cas9s. All engineered Cas9s were less promiscuous than wild type when canonical gRNA was used but HypaCas9 had much-reduced on-target unwinding. Cas9-HF1 and eCas9 showed the best balance between low promiscuity and high on-target activity with canonical gRNA. When extended gRNAs with one or two guanines added were used, Sniper1-Cas9 showed the lowest promiscuity while maintaining high on-target activity. Truncated gRNA generally reduced unwinding and adding a non-matching guanine to the 5’ end of gRNA influenced unwinding in a sequence-context dependent manner. Our results are consistent with cell-based cleavage data and provide a mechanistic understanding of how various Cas9/gRNA combinations perform in genome engineering.

## INTRODUCTION

CRISPR enzymes complexed with programmable guide-RNA (gRNA) can target complementary sequences in DNA or RNA, and their applications have revolutionized biology^1,2^. One of the most widely used CRISPR enzymes is Cas9 (SpCas9 from *Streptococcus pyogenes*) which in complex with a gRNA (Cas9-RNA) binds 20 base-pair (bp) long complementary DNA sequences (protospacer) that follow a motif called the protospacer adjacent motif (PAM; 5’-NGG-3’ for SpCas9) to form Cas9-RNA-DNA^3,4^. This binding involves unwinding of the protospacer and concomitant hybridization between gRNA and protospacer target strand (gRNA-DNA base-pairing)^3–8^. Following binding, Cas9-RNA activates its two nuclease domains (HNH for target strand and RuvC for non-target strand) and cleaves the DNA, producing a double-strand break^3–9^. Several Cas9 variants have been engineered (Engineered Cas9; EngCas9) to improve cleavage specificity^10–17^. Stable binding by Cas9s requires as few as 9-10 bp between the PAM-proximal end of the protospacer and the gRNA, but cleavage requires more bps (~16 for WT Cas9 and ~17-18 for EngCas9s)^5,7,18^. We previously showed, using a single molecule FRET unwinding assay, (Fig. 1a) that the DNA in Cas9-RNA-DNA undergoes internal unwinding-rewinding dynamics and the unwound state is a critical checkpoint required for cleavage. The intrinsic rate of cleavage (*k*_c, int_) has been used to refer to the rate of cleavage from the unwound state ^15,18^. The PAM-distal mismatches reduce the lifetime and extent of the unwound state, thereby delaying or preventing cleavage^18^. EngCas9s; eCas9^10^, and Cas9-HF1^19^, have higher cleavage specificity in part because their mutations make it difficult to unwind DNA when the sequence match is not perfect^18^. These variants also lower *k*_c, int_ thereby making it less likely that cleavage can occur from transiently unwound states^18^. Another strategy shown to improve specificity in genome engineering is the use of truncated^20^ and extended gRNAs^21,22^, collectively referred to as non-canonical gRNAs. However, the mechanisms underlying their specificity enhancements are not well understood. Here we have employed a smFRET assay to investigate the effects of non-canonical gRNAs^20–22^ and two newer EngCas9s, Hyper Accurate Cas9^12^ (HypaCas9; N692A/M694A/Q695A/H698A mutations) and SniperCas9^15^ (F539S/K890N/M763I mutations), on DNA unwinding (Supplementary Fig. 1). Because cell-based cleavage data are already available for most of these combinations of EngCas9s and non-canonical gRNAs^15^, we were able to compare on-target activities and specificity in unwinding to those of cleavage.

**Figure 1.**
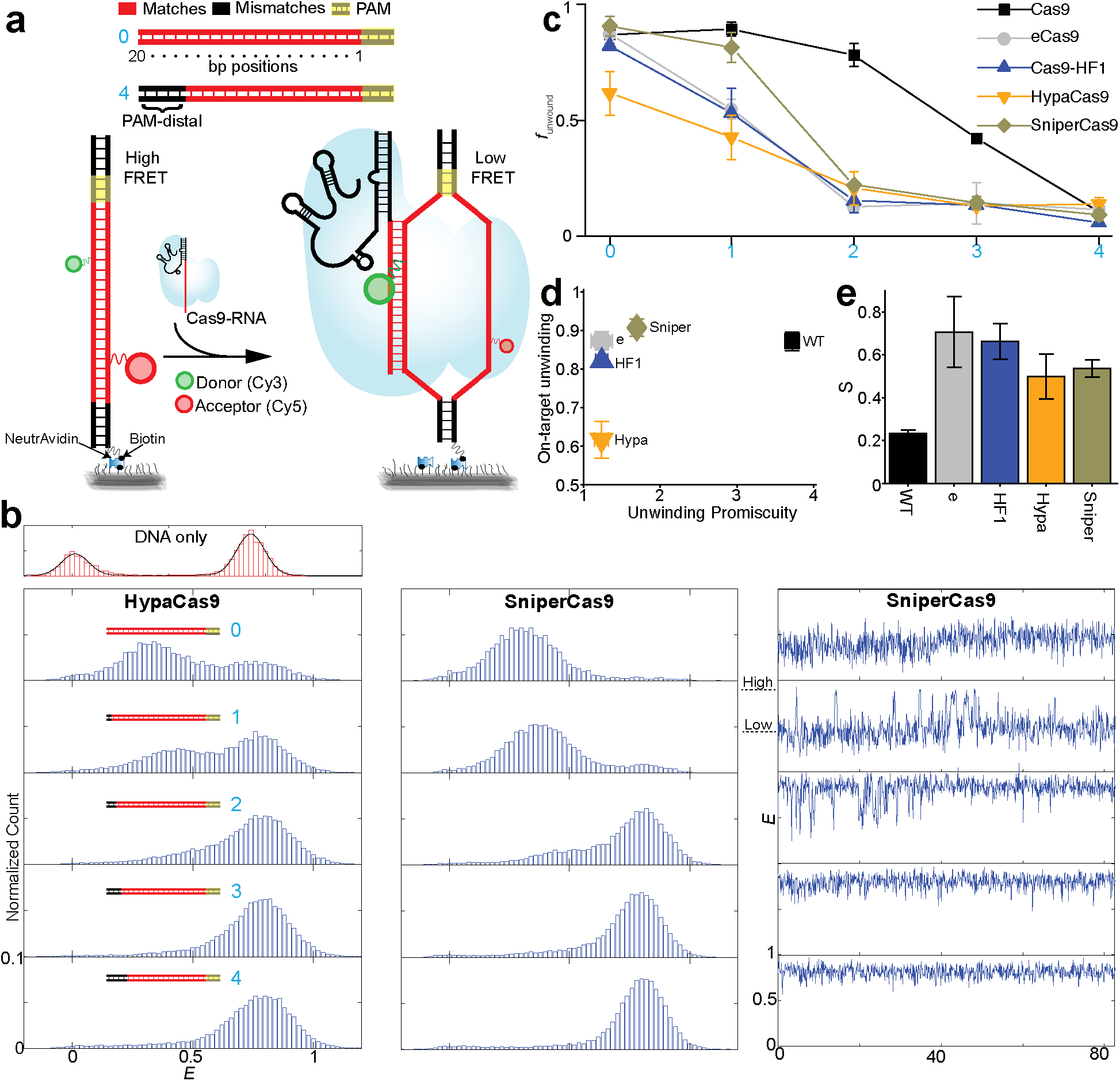
Unwinding specificity of HypaCas9 and SniperCas9. **(a)** Schematic of a smFRET assay for investigation of Cas9-RNA induced DNA unwinding of surface immobilized DNA incubated against free Cas9-RNA in solution. Cognate DNA and DNA with mismatches (no complementarity) against the gRNA at the indicated PAM-distal site in the protospacer region were used in this assay. The number of PAM-distal (*n*_PD_) mismatches is shown in cyan digits. **(b)** *E* histograms for DNA with different *n*_PD_ = 0, 1, 2, 3 and 4 (cyan digits) for dHypaCas9 and dSniperCas9. Also accompanied are representative *E* time-traces for dSniperCas9. **(c)** *f*_unwound_ vs. *n*_PD_ for different Cas9s. **(d)** A plot of on-target unwinding activity (*f*_unwound_ of cognate DNA) vs. unwinding promiscuity for different Cas9s. **(e)** Unwinding specificity (*S*_unwinding_) for different Cas9s. Error bars represent standard deviation (s.d.) from n=2. Data for dCas9, deCas9, and dCas9-HF1 is taken from a previous study^31^.

## Results

We previously developed a smFRET assay to investigate DNA unwinding in the PAM-distal region of the protospacer^23,24^. The DNA target molecules, with donor and an acceptor labels on the target and non-target strand, respectively, separated by 9 bp, were immobilized on a polymer-passivated quartz slide through biotin-neutravidin interaction. Without Cas9-gRNA, the FRET efficiency value (*E*) is ~0.75. Upon addition of Cas9-gRNA and subsequent DNA binding and unwinding, the distance between donor and acceptor increases, lowering *E* values (Fig. 1a and Supplementary Fig. 2). We used catalytically inactive versions for Cas9 (referred to as dCas9s) to avoid potential complications arising from DNA cleavage and will omit the prefix ‘d’ here. DNA targets are denoted by their number of PAM-distal mismatches relative to the gRNA (*n*_PD_). For example, *n*_PD_ is 2 for DNA with the 19^th^ and 2^th^ base pairs (counting from PAM) mismatched relative to the gRNA. We showed previously that for WT Cas9, the *E* value for cognate DNA (*n*_PD_=0) is reduced from 0.75 to 0.3 due to Cas9-induced unwinding^18^ and that as more PAM-distal mismatches are introduced, i.e. with increasing *n*_PD_, the low FRET unwound state is progressively converted to high FRET rewound state^18^ where about 9 PAM-proximal bps are unwound^25^. Further, we showed that the relative fraction of the unwound state, *f*_unwound_, generally decreases with increasing *n*_PD_, correlating with a decrease in the apparent cleavage rate, indicating that the unwound state is an obligatory intermediate for cleavage^18^. *f*_unwound_ vs. *n*_PD_ curves showed more rapid drops for eCas9 and Cas9-HF1 compared to WT Cas9 (Fig. 1b), and such enhanced sensitivity to PAM-distal mismatches contribute to their improved cleavage specificity^18^.

**Figure 2.**
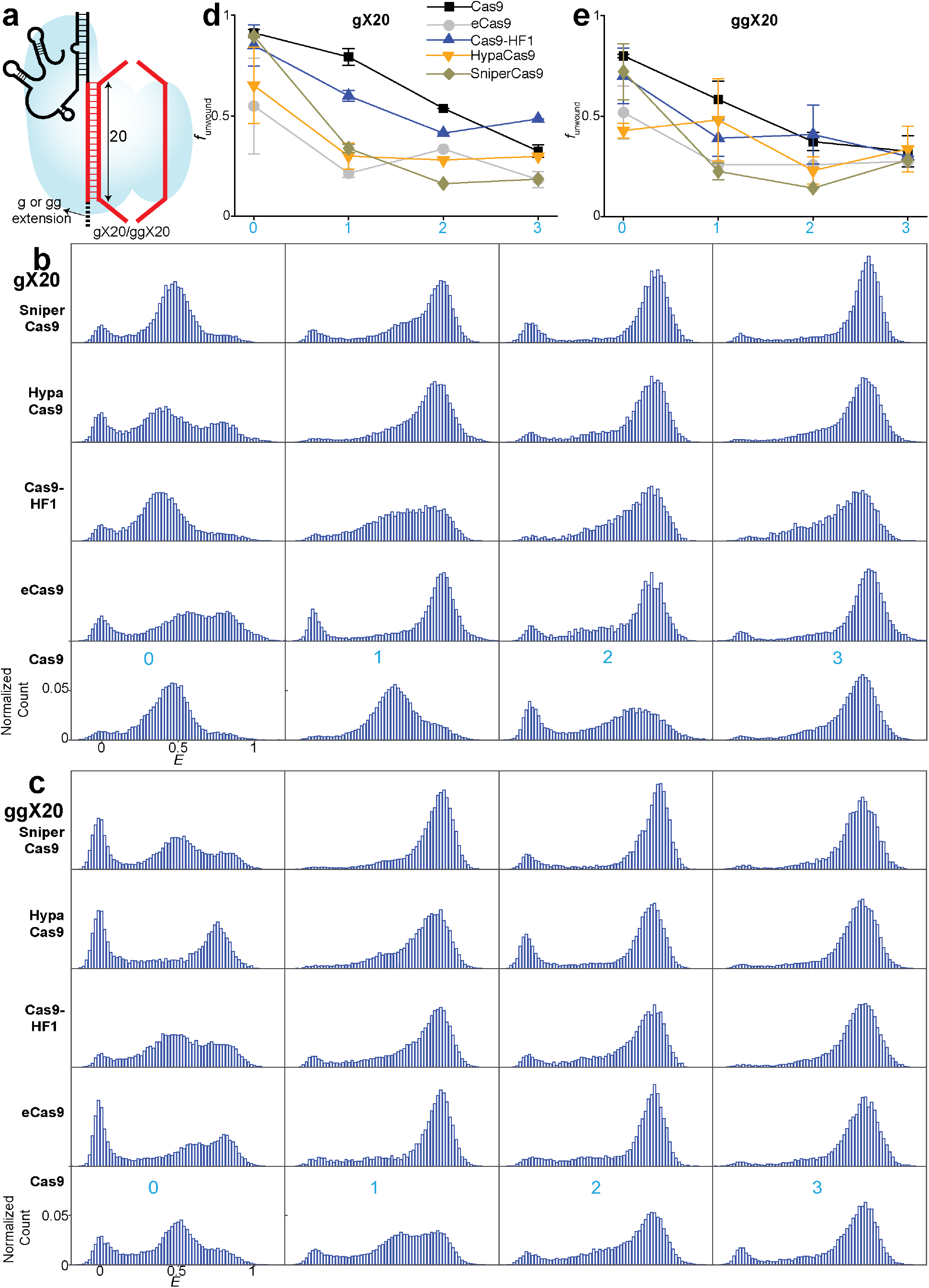
Unwinding by extended gRNA. **(a)** Schematic and naming convention of extended gRNA used in the unwinding assay. gX20 or ggX20 are gRNA with 5’g and 5’ gg non-hybridizing extensions respectively with canonical lengths of 20 bp in target strand hybridizing region. **(b-c)** *E* histograms of unwinding by extended gRNA. Each row corresponds to a particular labeled Cas9. Each column corresponds to a particular DNA target. **(b)** Histograms by gX20. **(c)** Histograms by ggX20. **(d)** *f*_unwound_ vs. *n*_PD_ for different Cas9s with gX20. **(e)** *f*_unwound_ vs. *n*_PD_ for different Cas9s with ggX20.

### HypaCas9 and SniperCas9 with canonical gRNA

First, we performed smFRET-based unwinding experiments on HypaCas9 and SniperCas9 with canonical gRNA. HypaCas9 and SinperCas9 also show the unwound and rewound states, with rewinding favored with increasing *n*_PD_(Fig. 1b-c). The two states interconverted within single molecules (Fig. 1b). Interestingly, HypaCas9 and SniperCas9 had weaker *n*_PD_dependence of *f*_unwound_ compared to eCas9 and Cas9-HF1 (but stronger dependence than WT Cas9), suggesting that their unwinding activities are more promiscuous compared to eCas9 or Cas9-HF1 (Fig. 1b-c).

For quantitative comparison of unwinding promiscuity across different variants of Cas9 and gRNA, we defined unwinding promiscuity as 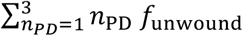. In this definition, if *f*_unwound_ does not decrease rapidly with increasing *n*_PD_, unwinding promiscuity is high with the penalty being higher for higher *n*_PD_. Figure 1d shows on-target unwinding activity [*f*_unwound_ (*n*_PD_=0)] vs. unwinding promiscuity for all five Cas9 variants with canonical gRNA. All engineered Cas9s display much-reduced unwinding promiscuity compared to WT Cas9 and their on-target unwinding activities are not impaired except for HypaCas9, which showed a lower on-target unwinding activity (Fig. 1d).

We also defined a single figure of merit, *S*, as *f*_unwound_ (0) / [unwinding promiscuity]. Ideally, what is needed is a system with high on-target activity and low promiscuity resulting in a high *S* value. *S* was, in increasing order, 0.23 for WT Cas9, 0.50 for HypaCas9, 0.53 for SniperCas9, 0.66 for Cas9-HF1 and 0.71 for eCas9 (Fig. 1e). *S* is low for WT Cas9 due to its high promiscuity. EngCas9s have higher *S* than WT Cas9, mainly because of reduced unwinding promiscuity. HypaCas9 has lower *S* than other EngCas9s due to its reduced on-target unwinding activity. In contrast, SniperCas9 has higher on-target unwinding activity but is more promiscuous compared to other engineered Cas9s.

### Extended gRNA

Next, we examined whether a system with higher *S* may be created by further altering the on-target/off-target activities of Cas9 variants through the use of non-canonical gRNAs. The canonical gRNA has 20 nucleotides (nt) (referred to here as X20) that are complementary to the protospacer. It has been shown that adding extra nucleotides at the 5’ end of canonical gRNA (X20) alters Cas9’s activity^21,22^. We wondered if Cas9 variants might have higher S value with the extended gRNAs. Therefore, we performed smFRET unwinding experiments on all five Cas9 variants with two extended gRNAs, referred to here as gX20 and ggX20, where g and gg represent extra non-hybridizing guanines on the 5’ end (Fig. 2a and Supplementary Fig. 3a). Because 5’ g or 5’ gg sequence can make U6/T7 transcription more efficient^26^, if gX20 or ggX20 does not reduce *S*, it would expand the repertoire of sequences that can be targeted for genome engineering.

**Figure 3.**
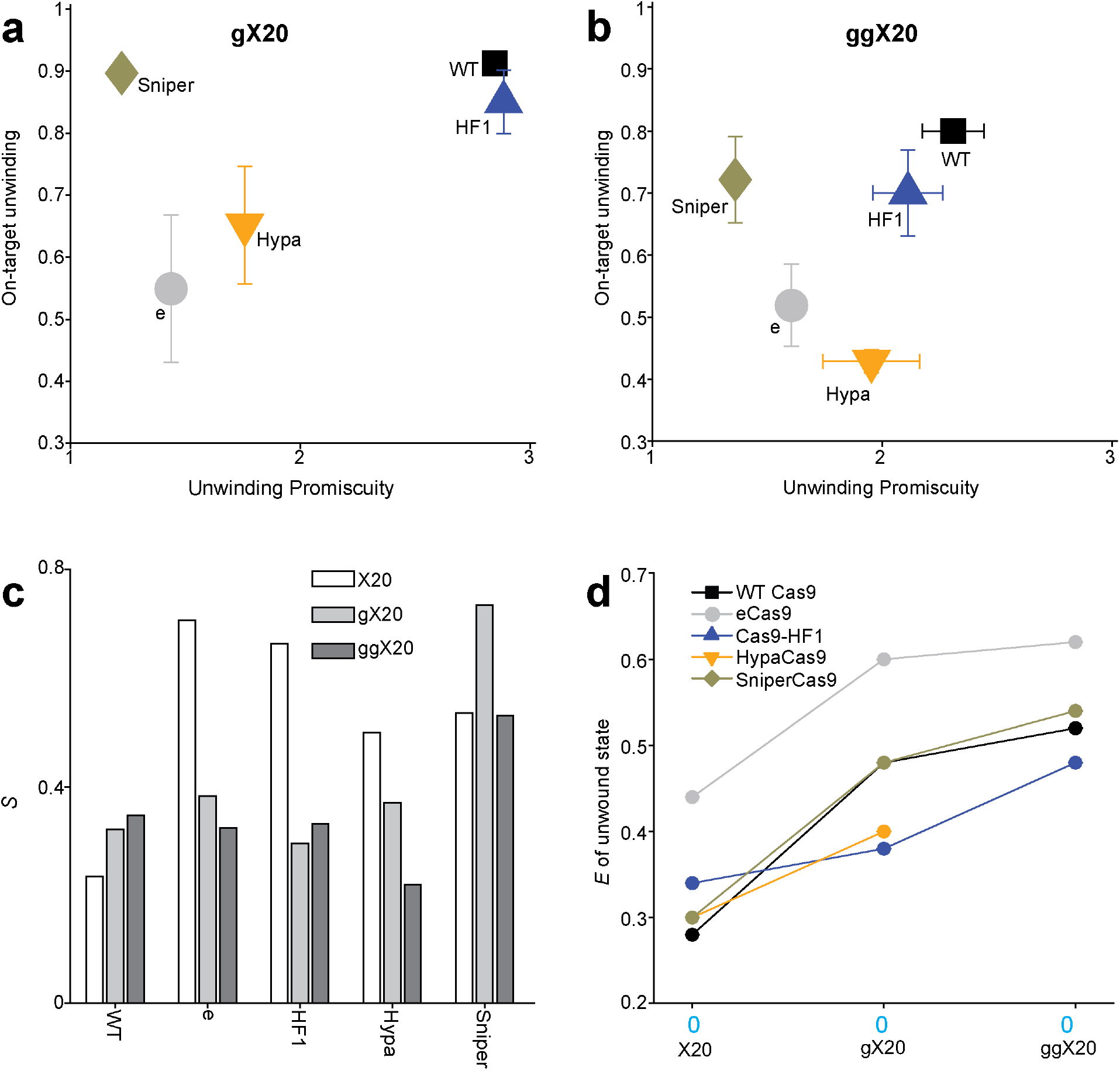
Unwinding specificity of extended gRNAs and the *E* values of their unwound state. **(a-b)** A plot of on-target unwinding activity vs. unwinding promiscuity with gX20 (a) and ggX20 (b). **(c)** *S* compared between canonical and extended gRNAs. **(d)** *E* values of the unwound state compared between canonical (X20) and extended gRNAs.

The unwound and rewound FRET states were also observed with gX20 and ggX20 (Fig. 2b-c), and *f*_unwound_ with different DNA targets were determined (Fig. 2d-e). SniperCas9 with gX20 had a much larger decrease in *f*_unwound_ with a single mismatch (*n*_PD_=1) than X20, and gX20 did not reduce on-target *f*_*unwound*_ (Fig. 2d and Fig. 3a-b). This result suggests that for SniperCas9, gX20 improves unwinding specificity while retaining the on-target unwinding activity of X20. Indeed, *S* values for SniperCas9 with gX20 and ggX20 were among the highest of all Cas9 and gRNA combinations tested, including those of canonical gRNAs (Fig. 3c). WT Cas9 also showed higher *S* values with the extended guide RNAs (Fig. 3c). In contrast, *S* was lower with extended gRNAs for eCas9, Cas9-HF1, and HypaCas9 (Fig. 3c). Therefore, SniperCas9 appears to be the best candidate for expanding the repertoire of sequences that can be targeted for genome engineering when a transcribed gRNA is used.

We also found *E* values of the unwound state increased with gX20 and increased further with ggX20 (Fig. 3d), suggesting that the extra guanines decrease the number of bp unwound. Finally, we note that, with gX20, Cas9-HF1 and HypaCas9 showed higher unwinding than eCas9 for most DNA targets (Fig. 2d).

### Truncated gRNA

It has been shown that the truncation of gRNAs at the 5’ end (PAM-distal side) by 2/3 nt, referred to here as X18/X17 respectively, also alters Cas9’s activity (Fig. 4a and Supplementary Fig. 3b)^20^. Therefore, we performed smFRET unwinding experiments using all five Cas9 variants with truncated gRNAs.

**Figure 4.**
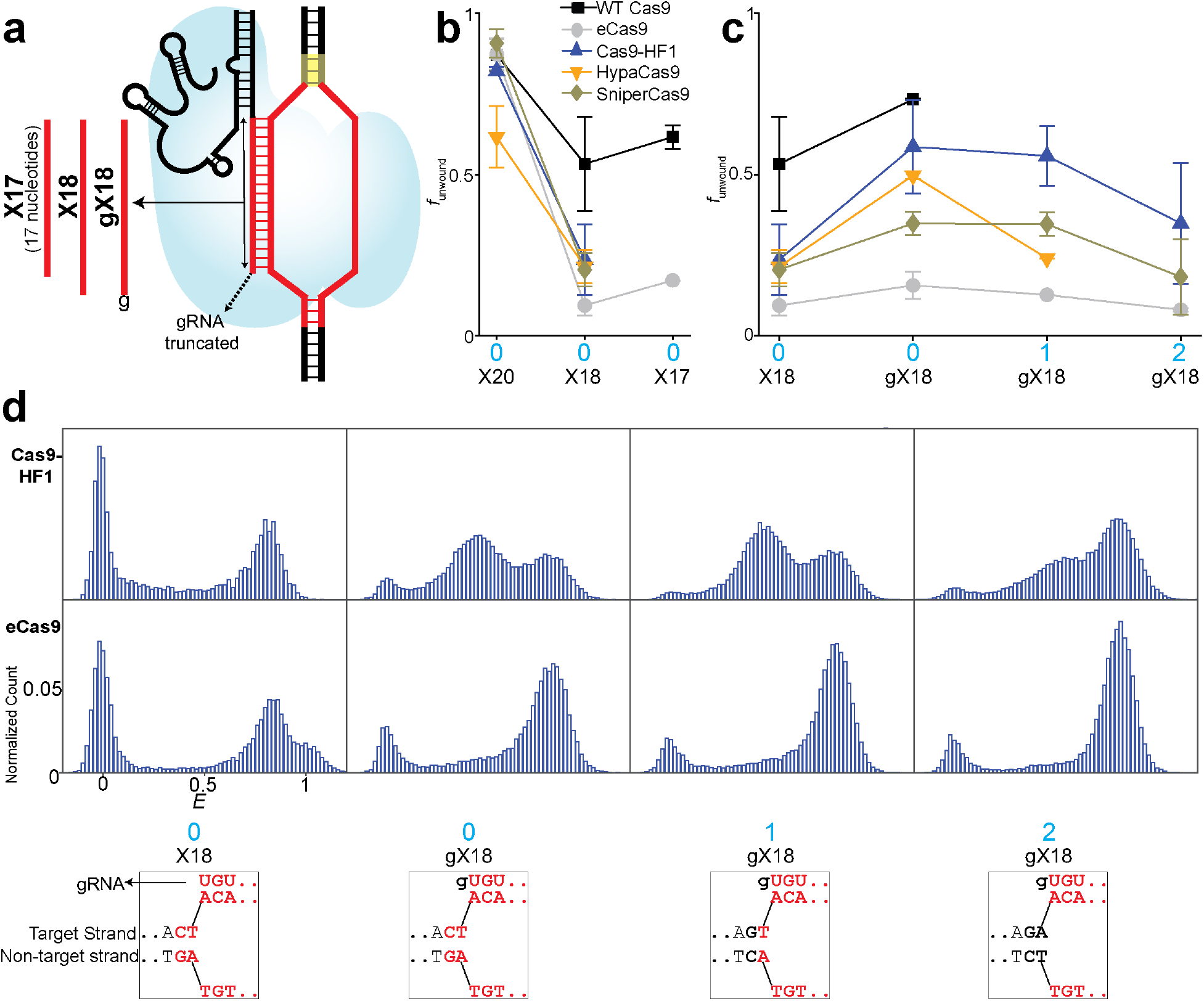
Unwinding by truncated gRNA. **(a)** Schematic and naming convention of truncated gRNA used in the unwinding assay. X18/X17 indicate the gRNA with length of 18/17 bp in the target strand hybridizing region, as opposed to the length of 20 bp for canonical gRNA. Addition of a mismatched g at the 5’ end of X18 gives gX18. **(b)** Comparison of *f*_unwound_ between X20, X18, and X17 for cognate DNA. **(c)** *f*_unwound_ with the different gRNA-Cas9 combinations for different DNA with 18 base pairs between gRNA and target strand. **(d)** *E* histograms of unwinding using truncated gRNA. The two rows indicate data for Cas9-HF1 and eCas9, respectively, whereas each column represents a particular DNA target and gRNA combination. The PAM-distal sequences of such gRNA and DNA combination are shown below for each column. All these DNA targets present the same number of 18 matching sequences to the gRNAs but differ in the sequence only beyond these 18 matching sequences at the PAM-distal site. Such differing nucleotides are marked in black. Data for dCas9, deCas9, and dCas9-HF1 with X20 is taken from a previous study^31^.

We observed both the unwound and rewound FRET states with X18 gRNA (**Supplementary Figure 4-5a**). *f*_unwound_ decreased significantly when X18/X17 replaced X20 (Fig. 4b). This decrease is expected because without full base pairing between gRNA and the target DNA strand, the unwound state would be more difficult to sustain. Adding an unmatched guanine to the 5’ end of X18 (referred to here as gX18) increased *f*_unwound_ suggesting that the extra nucleotides affect unwinding even though it does not base pair with the target strand (Fig. 4c-d and Supplementary Figure 5). In addition, using gX18 against three DNA targets having the same 18 bp matches but different nucleotides in the 19^th^ and 20^th^ positions, counting from the PAM, showed different *f*_unwound_ values (Supplementary Figure 5). Overall, the data suggest that unwinding depends on extra 5’ nucleotides added to X18 and also mildly on the DNA sequences of PAM-distal region. Because truncated gRNAs cannot base pair with the full 20 nt of the protospacer, unwinding promiscuity or *S* values cannot be defined.

As was the case with an extended gRNA gX20, Cas9-HF1 and HypaCas9 consistently showed higher unwinding than eCas9 also with gX18 (Fig. 4c-d). This difference in unwinding activity between Cas9-HF1/HypaCas9 and eCas9 with gX20 and gX18 is remarkable because no such difference was observed amongst EngCas9s with a canonical gRNA (Fig. 1c, Fig. 2b, Fig. 2d and Fig. 4c-d).

## Discussion

HypaCas9 mutations were rationally engineered into the Cas9’s REC3 domain which recognizes the unwound state and transmits the signal to HNH for nuclease activation^8,12,18^. The reduced unwinding of the cognate DNA target for HypaCas9 suggests that contacts between the target strand and the polar residues of REC3, mutated in HypaCas9, stabilize the unwound state. Despite a more rational approach behind the design of HypaCas9^12^, its on/off target ratio in cellular cleavage efficiency is not notably different from eCas9/Cas9-HF1^15^, potentially due to its low on-target unwinding activity we observed here.

Structures of Cas9-gRNA-DNA with X20 show that the 5’ end of the gRNA makes direct contacts with the RuvC domain of Cas9 ^27 26^ (Supplementary Figure 6a), effectively capping the 5’ end of the gRNA. Therefore, accommodation of extensions on the 5’ end of the gRNA is likely to cause distortion of the rRNA-DNA, and thus the destabilization of the unwound state, as seen in the structure of Cas9-gRNA-DNA with ggX20 (Supplementary Fig. 6a)^28^. Our observation that *E* values of the unwound state increase with gRNA extension is consistent with such structural distortion.

A previous study used smFRET analysis of doubly labeled WT Cas9 to monitor the movement of HNH domain between inactive and active states, and showed that the fraction of HNH activated state, *f*_HNH_, decreases with increasing *n*_PD_ and with gRNA truncation to 17 nt (X17)^29^, paralleling our observation of *f*_unwound_ decrease with increasing *n*_PD_ and gRNA truncation. For WTCas9 with X20, *f*_HNH_ is 78-100% of *f*_unwound_ for all DNA targets indicating that most unwound states lead to HNH activation with canonical gRNA^29^. *f*_HNH_ with X17 was 45% of *f*_unwound_ with X17^29^, indicating more than half of unwound states do not generate HNH activation with X17. Because REC3 senses DNA unwinding and concomitant formation of RNA-DNA hybrid via its interactions with both target strand and gRNA ^27,30^, gRNA truncation may make it difficult for REC3 to recognize the RNA-DNA hybrid and transmit DNA unwinding towards downstream HNH activation. EngCas9s generally showed reduction in unwinding, HNH activation, and intrinsic rate of cleavage from unwound state^12,31^. Their combination with truncated gRNA diminishes their unwinding further (Fig. 4b-c and Supplementary Fig. 4), in part accounting for EngCas9s’ low on-target cleavage activities with truncated gRNA in cells, except SniperCas9 which retained some activity^11,15^.

The unmatched guanine(s) of gX18/gX20/ggX20 may form interactions with the DNA target, as observed in the structure of Cas9-ggX20-DNA^32^ (Supplementary Fig. 6b). Such interactions may potentially influence unwinding activity in a manner dependent on PAM-distal DNA sequence as we observed with gX18. In the Cas9-ggX20-DNA structure, the RNA-DNA hybrid is repositioned relative to the Cas9 residues that it interacts with when the canonical gRNA is used (Supplementary Fig. 6b-c). Because Cas9-HF1 and HypaCas9 reduce unwinding through mutations to these residues, we speculate that repositioning of the hybrid induced by the extra guanine(s) partially relieves DNA unwinding defects of Cas9-HF1/HypaCas9 mutations. This partial relief leads to significant increases in Cas9-HF1/HypaCas9’s unwinding compared to eCas9, whose mutations affect non-target strand and not the RNA-DNA hybrid.

In some cases, WT Cas9 combined with non-canonical gRNA may improve specificity while maintaining high on-target activity. In addition, certain EngCas9s may perform the best when coupled with non-canonical gRNA. For example, SniperCas9 is the only EngCas9 with significant cleavage activity in cells with both the extended and truncated gRNA^15^. Although SniperCas9 is the most promiscuous in unwinding among all EngCas9s when the canonical gRNA is used (Fig. 1b-e), it showed the largest discrimination against a PAM-distal single base pair mismatch amongst all Cas9 and gRNA combinations tested when combined with gX20. This result potentially explains why the SniperCas9/gX20 combination produced the largest on-target/off-target cleavage activity in cells ^15^.

The types of non-canonical gRNA are also continually expanding, including gRNA with DNA nucleotides^33^, chemical modifications^34,35^ whose specificity improvement may also arise from DNA unwinding becoming more sensitive to mismatches. Non-canonical gRNAs can expand genome engineering applications a few additional ways. First, efficient T7 or U6 transcription of gRNA require 5’g or 5’gg^26,36^. Due to this requirement, only the gRNA with 5’g in its target strand hybridizing region are routinely employed thus restricting the number of available sites^37^. However, the 5’g and 5’gg extensions in the gRNA not only satisfies this requirement to expand potential target sites but also provides improved specificity for wild type Cas9 and SniperCas9. Second, transfection of non-canonical gRNA may provide an alternative path toward obtaining higher specificity in existing cell and animal systems that express WT Cas9. Third, the ultrastability of Cas9-RNA-DNA even after cleavage may prevent the exposure of the cleaved site until motor proteins such as RNA polymerases can help remove Cas9-RNA from the DNA^38^. Weakening the Cas9-RNA-DNA complex by EngCas9 mutations^39^ or non-canonical gRNA may facilitate Cas9-RNA removal and gene editing. Therefore, an appropriate combination of EngCas9 and/or non-canonical gRNA may provide not only an improved on-target cleavage activity and specificity but also higher gene editing efficiencies.

## ONLINE METHODS

### Preparation of DNA targets

The schematic of the DNA target used in the smFRET assay to investigate Cas9-RNA induced DNA unwinding is shown in Supplementary Figure 2–3. Integrated DNA Technologies (IDT, Coralville, IA 52241) was the commercial supplier of all DNA oligonucleotides. For introducing Cy3 and Cy5 labels at the indicated locations, the oligonucleotides were purchased with amine-containing modified thymine at the indicated location. A C6 linker (amino-dT) was used to label the DNA strands with Cy3 or Cy5 N-hydroxysuccinimido (NHS). For preparing the DNA, the constituents non-target strand, target strand, and a 22 nt biotinylated adaptor strand were first mixed in 10 mM Tris-HCl, pH 8 and 50 mM NaCl. The mixture was then transferred to a heat block preheated to 90 °C. After 2 minutes of heating, the mixture was taken off the heat block and allowed to cool to room temperature over a few hrs. The sequences of the target and non-target strand (with same label positions) were changed to create DNA targets with mismatches. The full sequence of all DNA targets used in the smFRET assay is shown in Supplementary Table 1.

### Expression and purification of Cas9

The protocols for Cas9 expression and purification have been described previously^3,5,18^. pET-based expression vectors were used for expression of all Cas9s except SniperCas9 and dSniperCas9. For WT Cas9, the vector consisted of sequence encoding Cas9 (1-1368 residues of Cas9 from *Streptococcus pyogenes*) and an N-terminal decahistidine-maltose binding protein (His10-MBP) tag, followed by a peptide sequence containing a tobacco etch virus (TEV) protease cleavage site. This vector was used as a base vector for other Cas9s. The mutations for catalytically inactive cas9 (dCas9) and/or EngCas9 mutations were introduced by site-directed mutagenesis kits (QuickChange Lightning; Agilent Technologies, Santa Clara, CA 95050). Around-the-horn PCR was performed on the base vector for creating HypaCas9 mutations.

The cells were grown in TB (Terrific Broth) or 2YT medium (higher expression obtained for TB) at 37 °C for a few hours and the cell density was continuously monitored by measuring the optical density of the media at 600 nm (OD_600_). Cas9 expression was induced when the OD_600_ reached 0.6 with 0.5 mM IPTG, and the temperature was lowered to 18 °C. The cells were induced for 12-16 hours. Following which, the media containing cells were centrifuged to harvest the cells. The supernatant media, devoid of cells, was discarded. The harvested cells were then collected and lysed in the buffered solution containing 50 mM Tris pH 7.5, 500 mM NaCl, 5% glycerol, 1 mM TCEP, protease inhibitor cocktail (Roche) and with/without Lysozyme (Sigma Aldrich). Fisher Model 500 Sonic Dismembrator (Thermo Fisher Scientific) was used to homogenize the lysed cells by operating it at 30% amplitude in 3 one-minute cycles, each consisting of series of 2 s sonicate-2 s repetitions, or the lysed cells were homogenized in a microfluidizer (Avestin). The solution was subjected to ultra-centrifugation at 15,000 *g* for 30-45 minutes to remove the cellular debris from the lysed and homogenized solution. Following which, the supernatant of lysate was collected, and cellular debris was discarded.

The supernatant was added to Ni-NTA agarose resin (Qiagen), and His10-MBP-TEV-Cas9 was allowed to bind Ni-NTA. The Ni-NTA bound with His10-MBP-TEV-Cas9 was then washed extensively with 50 mM Tris pH 7.5, 500 mM NaCl, 10 mM imidazole, 5% glycerol, 1 mM TCEP. The His10-MBP-TEV-Cas9 bound to Ni-NTA was then eluted in a single-step with the elution buffer of 50 mM Tris pH 7.5, 500 mM NaCl, 300 mM imidazole, 5% glycerol, 1 mM TCEP. For buffer exchange and to remove imidazole from the eluted solution containing Cas9, the eluted solution was dialyzed into 20 mM Tris-Cl pH 7.5, 125 mM KCl, 5% glycerol, 1 mM TCEP) overnight at 4 °C. Concomitant with this dialysis, the cleavage of TEV-protease site by TEV protease was simultaneously carried out which resulted in free 10-His-MBP and Cas9 constituents in the solution. To remove and arrest away 10-His-MBP from the solution, the solution containing free 10-His-MBP and Cas9 constituents was subjected to another round of Ni-NTA agarose column purification resulting in the solution containing only Cas9. For further purification, this solution was subjected to size-exclusion chromatography on a Superdex 200 16/60 column (GE Healthcare) in Cas9 storage buffer (20 mM Tris-Cl pH 7.5, 100 mM KCl, 5 % glycerol and 5 mM MgCl_2_).

In some preparations, TEV cleavage and dialysis were not concomitant. The eluted solution containing His10-MBP-TEV-Cas9 fusion was first subjected to TEV protease treatment over a few hours. After TEV cleavage, the solution was then dialyzed into 20 mM Tris-Cl pH 7.5, 125 mM KCl, 5% glycerol, 1 mM TCEP for 3 hours. The dialyzed solution was then applied to HiTrap SP HP sepharose column (GE Healthcare) and washed with 20 mM Tris-Cl pH 7.5, 125 mM KCl, 5% glycerol, 1 mM TCEP for 3 column volumes. The Cas9 bound to the column was then eluted into 20 mM Tris-Cl pH 7.5, 1 M KCl, 5% glycerol, 1 mM TCEP using a linear gradient from 0-100% over 20 column volumes. The eluted Cas9 was then further purified into Cas9 Storage Buffer (20 mM Tris-Cl pH 7.5, 200 mM KCl, 5% glycerol, 1 mM TCEP) by size exclusion chromatography on a Superdex 200 16/60 column (GE Healthcare). All the purification steps described in this section were performed at 4 °C, and the purified Cas9 was stored at −80 °C for long-term storage. The SniperCas9 was also expressed and purified from *E. coli*. The DNA sequence encoding SniperCas9 along with an NLS, HA epitope and His-tag at the N-terminus was cloned into pET28-b(+) vector. Using this vector, the recombinant protein was then expressed in *E. coli*, and first purified from the harvested cell lysate using Ni-NTA agarose beads (Qiagen). For buffer exchange, this purified protein was then dialyzed against 20 mM HEPES pH 7.5, 150 mM KCl, 1 mM DTT, and 10% glycerol. The Ultracel 100 K cellulose column (Millipore) was then used to concentrate the purified/dialyzed SniperCas9. The SDS-PAGE was used to analyze the concentration and purity of the final SniperCas9 protein.

### Preparation of gRNA and Cas9-gRNA

The canonical gRNA of Cas9 consists of CRISPR RNA (crRNA) and trans-activating crRNA (tracrRNA) (Supplementary Figure 2) where the gRNA sequence hybridizing with the target strand of DNA target is 20 bp long^3,4^. All gRNAs were prepared by mixing crRNA and tracrRNA in 1:1.2 ratio in 10 mM Tris HCl (pH 8) and 50 mM NaCl. This mixture was then placed in a heating block pre-heated to 90 °C for 2 minutes. Following which, the mixture was allowed to cool to room temperature over a few hours for efficient hybridization between crRNA and tracrRNA. The non-canonical gRNA, i.e., truncated and extended gRNA was created by using the truncated and extended crRNA, respectively with the same tracrRNA (Supplementary Figure 3). Both the crRNA and tracrRNA were in-vitro transcribed as described previously and were purified to remove rNTPS using the commercial kits (Zymo Research). The in-vitro transcription produces RNA with 1-3 nt truncations and elongations at the 3’ end. Such likely truncations and elongations did not affect our experiments as the 3’ end of RNA have no role in Cas9-RNA activity, which was also confirmed by the control unwinding experiments using gel-purified fixed length RNAs which produced the same results as the kit-purified RNAs.Cas9-RNA was prepared by mixing the gRNA and Cas9 at a ratio of 1:3 in Cas9-RNA activity buffer of 20 mM Tris HCl (pH 8), 100 mM KCl, 5 mM MgCl_2_, 5% v/v glycerol. The full sequences of all the gRNA used in this study are available in Supplementary Table 1.

### Single-molecule fluorescence imaging and data analysis

Flow chamber surfaces coated with polyethylene glycol (PEG) was used for immobilization of DNA targets. These flow chambers were purchased from Johns Hopkins University Microscope Supplies Core. The neutrAvidin-biotin interaction was used for immobilizing the biotinylated DNA target molecules on the PEG-passivated flow chamber in the Cas9-RNA activity and imaging buffer (20 mM Tris-HCl, 100 mM KCl, 5 mM MgCl_2_, 5% (v/v) glycerol, 0.2 mg ml^−1^ BSA, 1 mg ml^−1^ glucose oxidase, 0.04 mg ml^−1^ catalase, 0.8% dextrose and saturated Trolox (>5 mM))^20^. Cas9-RNA in the Cas9-RNA activity and imaging buffer was added to the flow chamber at the concentrations much higher (E.g., 100 nM) than the dissociation constant of Cas9-RNA-DNA^18^ for Cas9-RNA targeting of DNA and Cas9-RNA induced DNA unwinding. All the imaging experiments were done at room temperature and time resolution was either 100 ms or 35 ms per frame. The total fluorescence from each of the immobilized DNA targets molecules was optically split into two separate donor and acceptor optical paths. The emissions belonging to these parts were projected onto two halves of a cryo-cooled (< −70 °C) EMCCD camera (Andor) which was stored as a video recording by the camera. The video recording containing fluorescent spots was then analyzed using custom scripts to extract background corrected donor fluorescence (I_D_), acceptor fluorescence (I_A_). FRET efficiency (*E*) of each detected spot was approximated as *E* = *I*_A_/(*I*_D_+*I*_A_). In the analysis of DNA unwinding experiments, the DNA molecules with the missing or inactive acceptor label were avoided by only including the fluorescent spots in the acceptor channel. Data acquisition and analysis software are available at https://cplc.illinois.edu/software/. Most of the primary analysis was carried out using custom-written scripts. Methods describing the acquisition of smFRET data and analysis have been reported previously^24^.

### *E* histograms and analysis of Cas9-RNA induced DNA unwinding and rewinding

For every single molecule, the first five data points of its *E* time-traces were used as data points to construct *E* histograms. More than 2000 molecules contributed to each *E* histogram. The donor only peak (*E*=0), low FRET (0.2 <*E* <0.6 or 0.65 or 0.70) and high FRET (*E* >0.6 or 0.65 or 0.7) are three characteristic populations observed in these *E* histograms. Based on this low and high FRET populations, Cas9-RNA induced DNA unwinding was modeled as a two-state system, as shown below. The unwound fraction (*f*_unwound_) was calculated as a fraction of the low FRET population in the *E* histograms of DNA unwinding experiments.

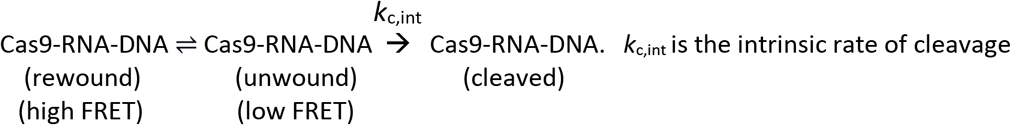

### Calculation of *S*

*S* was calculated as the ratio of *f*_*unwound*_ for cognate DNA and aggregate of *f*_*unwound*_ for DNA with PAM-distal mismatches, weighted by their *n*_*PD*_.

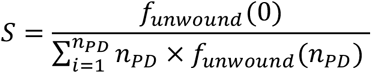

*n*_*PD*_ is the number of PAM-distal mismatches, *f*_*unwound*_ (0) is the unwound fraction for cognate DNA (*n*_*PD*_ = 0), and (*f*_unwound_= 0) and *f*_*unwound*_ (*n*_*PD*_) is the unwound fraction for DNA with a given *n*_*PD*_.

## ACKNOWLEDGMENTS

The project was supported by grants from the National Science Foundation (PHY-1430124 to T.H.) and the National Institutes of Health (GM 122569 to T.H. and GM 097330 to S.B.). T.H. is an investigator with the Howard Hughes Medical Institute.

---

**Supplementary Text Table 1:**
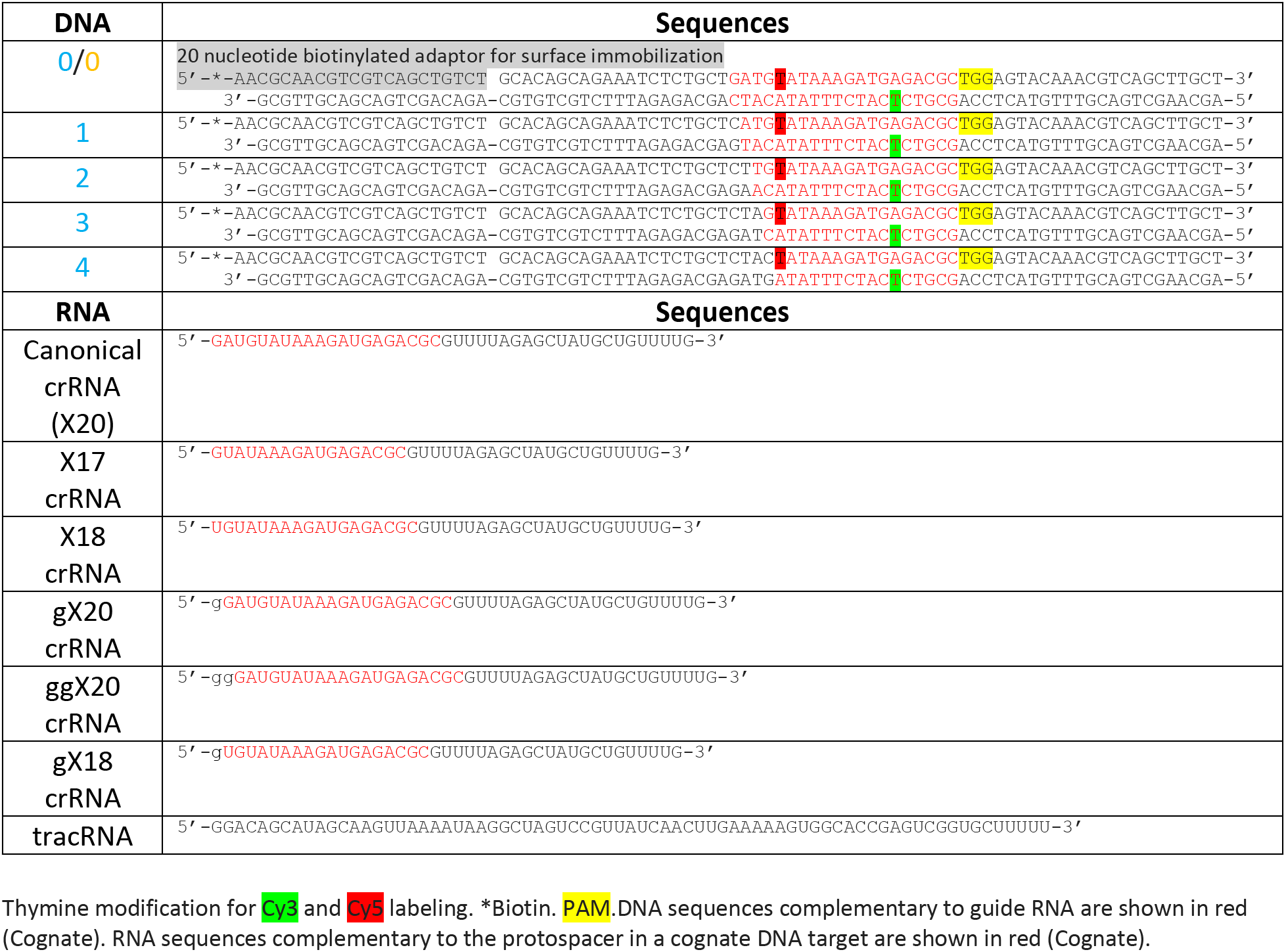
gRNA and DNA targets used for Cas9-RNA induced DNA unwinding smFRET assay.

## Supplementary Information

**Supplementary Figure 1.**
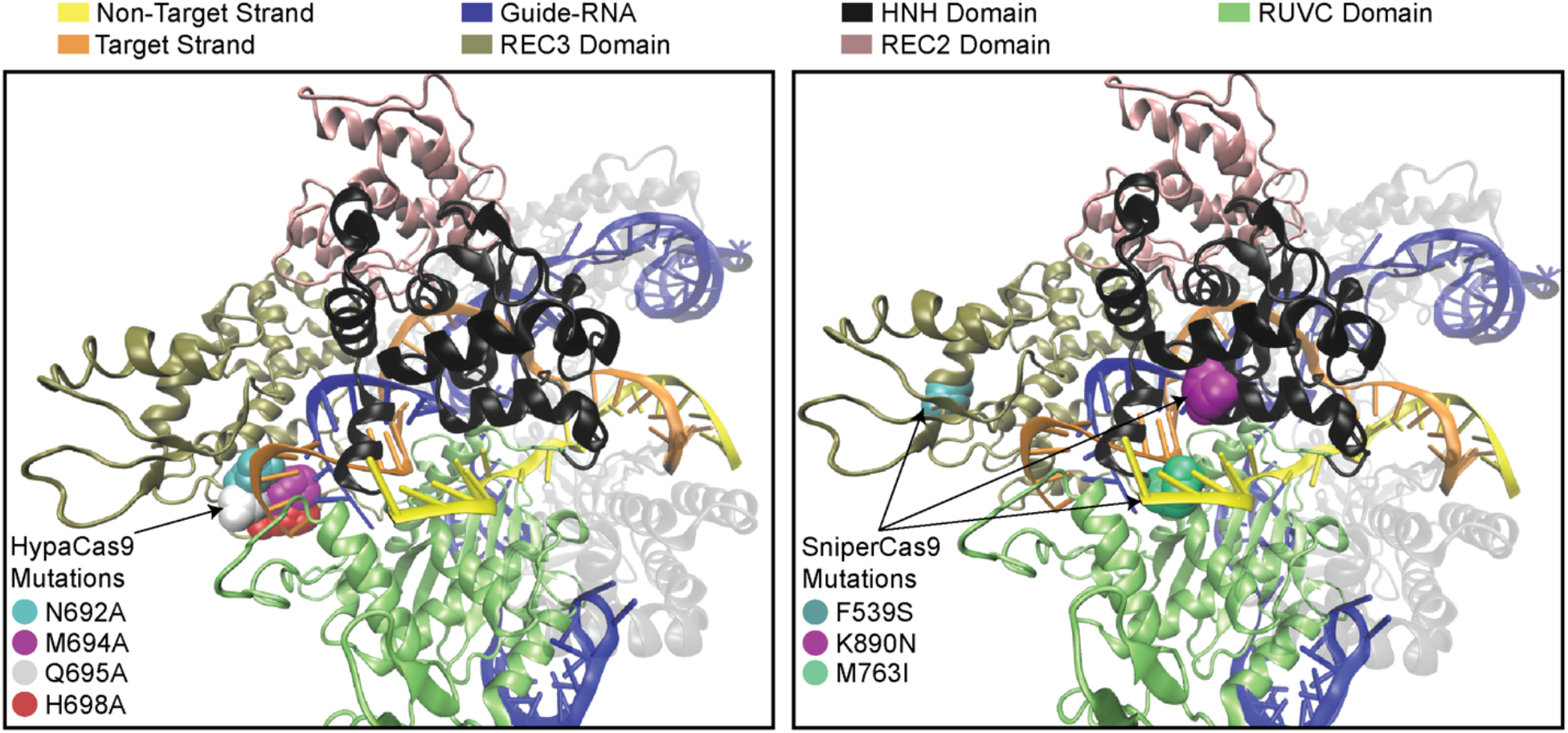
Locations of HypaCas9 and SniperCas9 mutations in the Cas9-RNA-DNA. (PDB ID: 5F9R^1^).

**Supplementary Figure 2.**
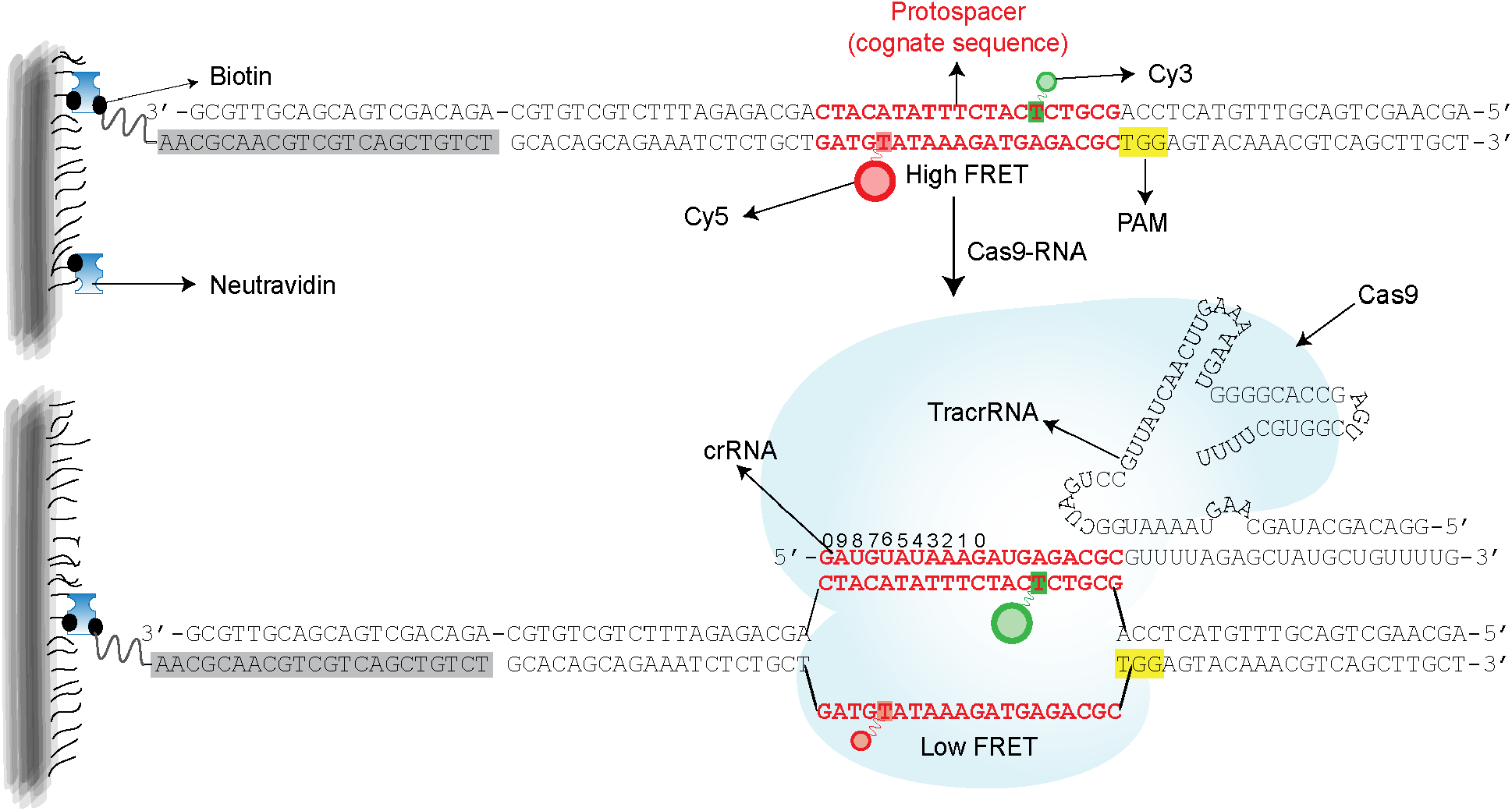
Locations of FRET probes for smFRET DNA unwinding experiments and DNA unwinding by WT Cas9 compared with EngCas9. Schematic of Cas9-RNA-DNA complex. crRNA hybridized with tracrRNA is the guide-RNA, and the denoted red sequences hybridize with one of the strands (target strand) in the DNA. The strand complementary to target strand in the DNA is the non-target strand which also contains the PAM (5’-NGG-3’) as indicated. Highlighted in grey is a 22 nt biotin-labeled strand which is used as an adaptor for surface immobilization of the DNA. The donor and acceptor labels are conjugated to the 6^th^ nucleotide (from PAM) in target strand and 16^th^ in the non-target strand. Cas9-RNA binds the DNA leading to the unwinding of the DNA which results in an increase in distance between FRET labels and thus a decrease in *E*.

**Supplementary Figure 3.**
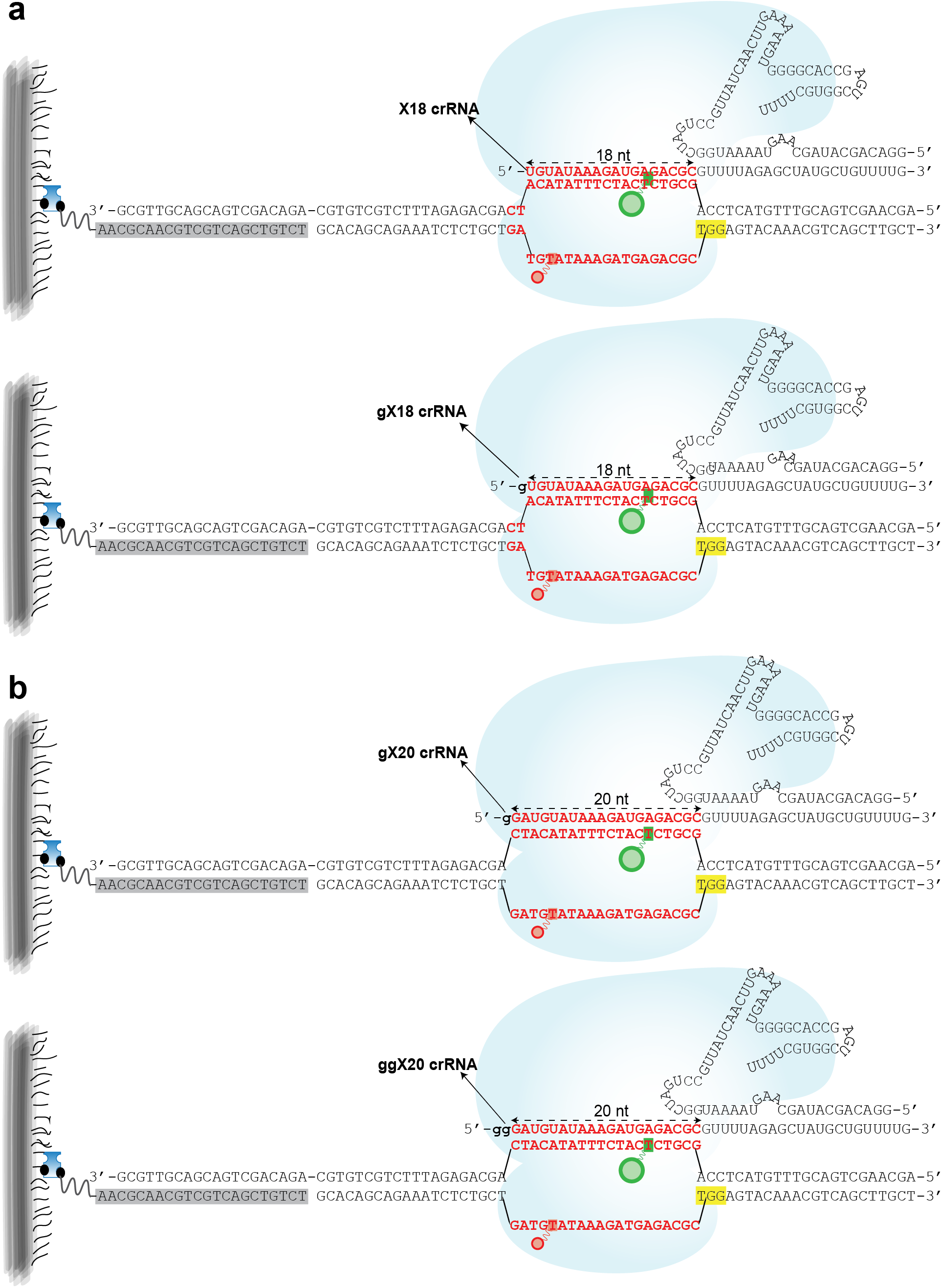
Schematic of the smFRET unwinding assay involving non-canonical gRNAs. (a) Extended gRNAs. (b) Truncated gRNAs.

**Supplementary Figure 4.**
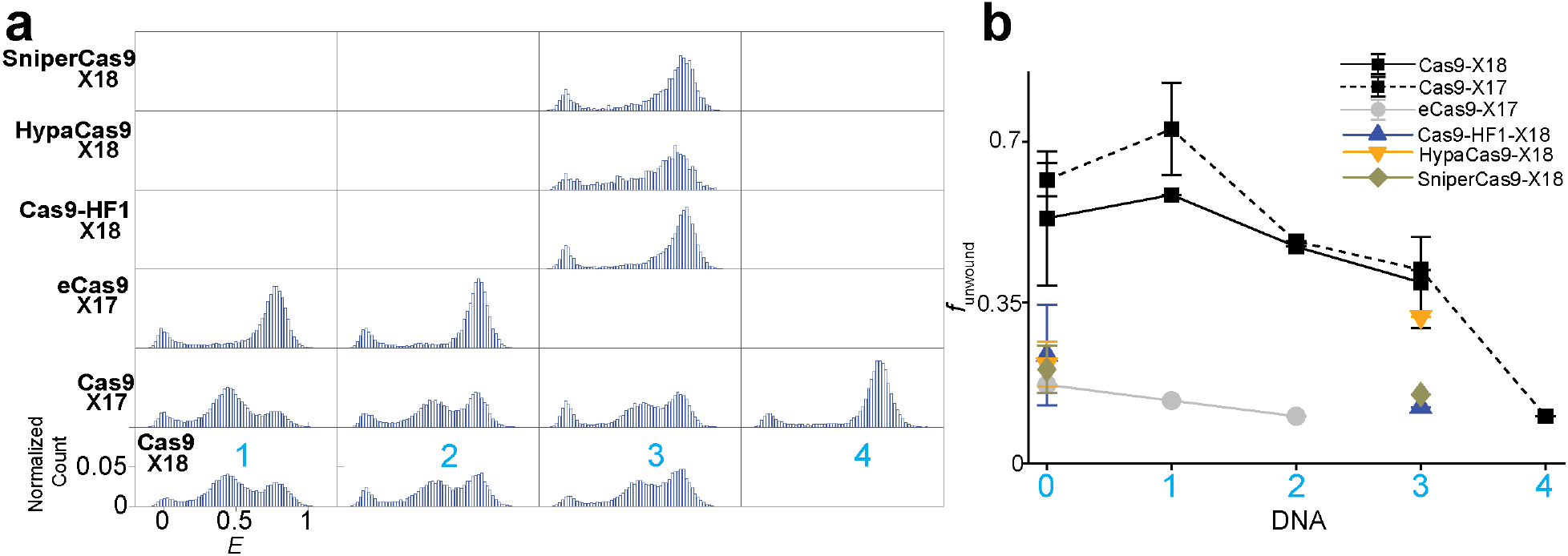
Unwinding by truncated gRNA. **(a)** *E* histograms of unwinding by truncated gRNA. Each row corresponds to a particular labeled Cas9 and gRNA combination. Each column corresponds to a particular DNA target. **(b)** *f*_unwound_ vs. *n*_PD_ for different Cas9s with truncated gRNAs.

**Supplementary Figure 5.**
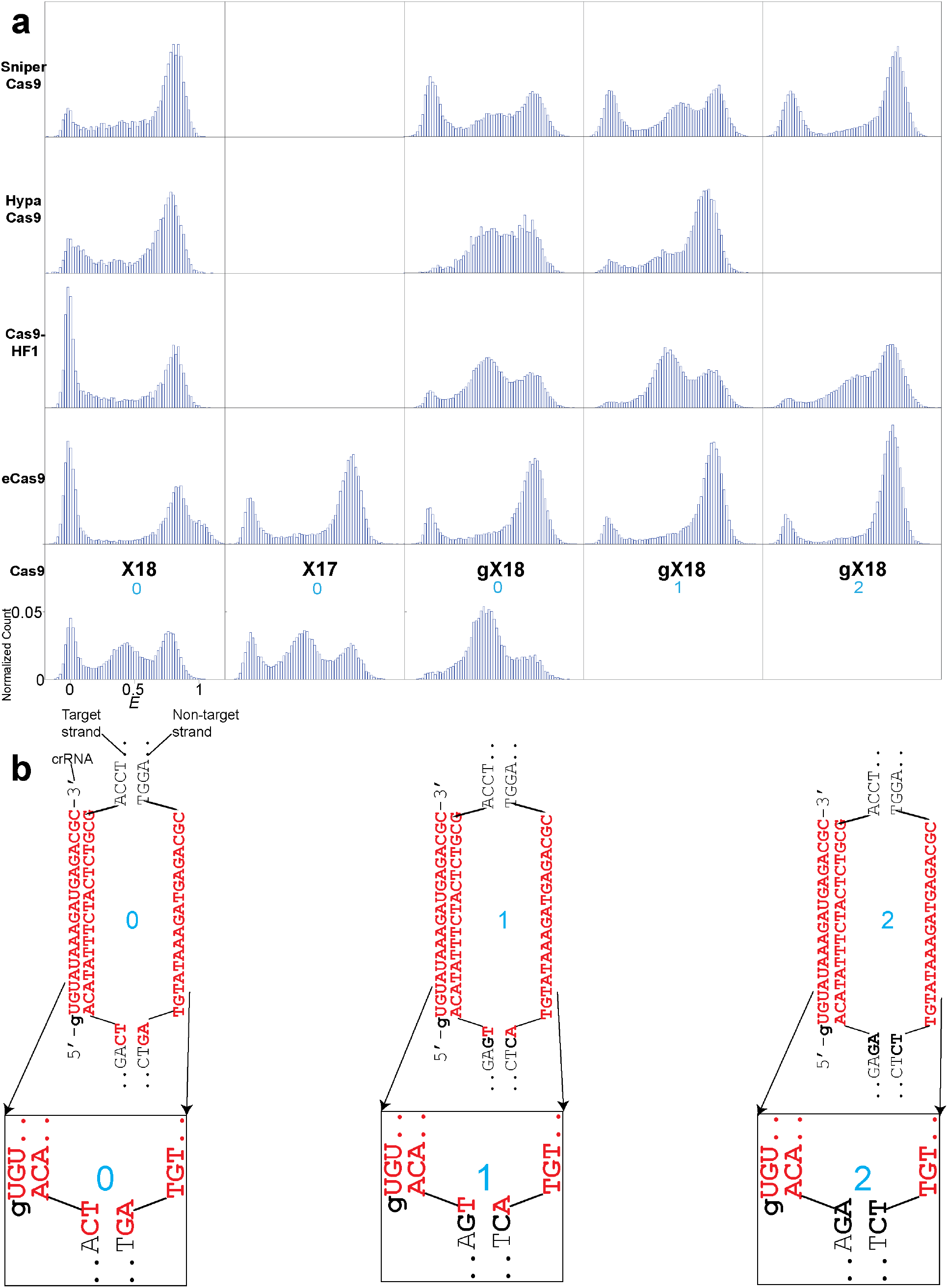
Unwinding by truncated gRNA. **(a)** *E* histograms of unwinding by truncated gRNA. Each row corresponds to a particular labeled Cas9. Each column corresponds to a particular gRNA and DNA target. **(b)** Sequences of DNA and crRNA (of truncated gRNAs). X18/gX18 guide RNA forms the same Watson-Crick base pairing with *n*_PD_=0, *n*_PD_=1, and *n*_PD_=2 DNA targets.

**Supplementary Figure 6.**
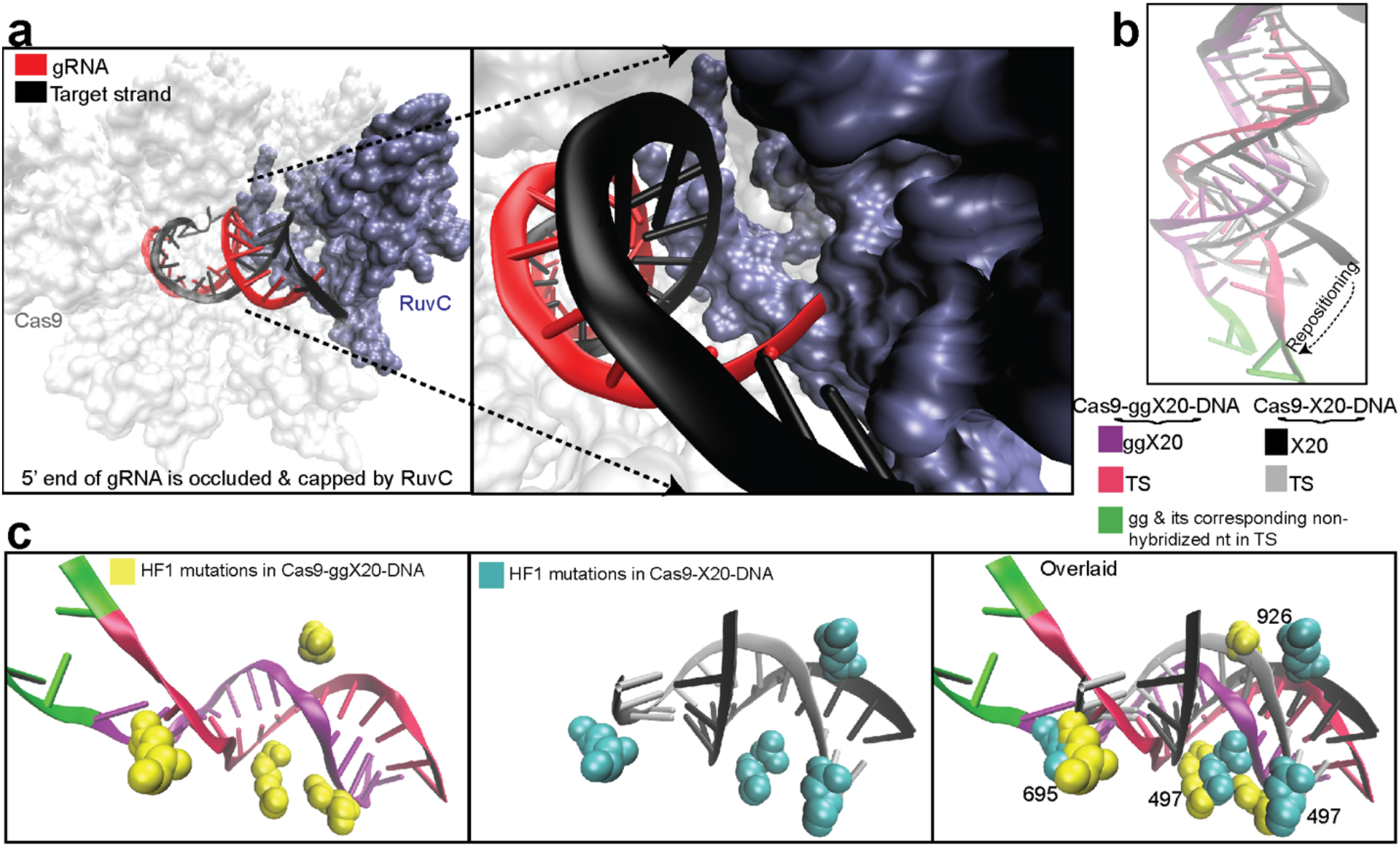
PAM distal end of Cas9-RNA-DNA complex. **(a)** 5’ end of the gRNA is occluded and capped by the RuvC nuclease domain in Cas9-X20-DNA. PDB ID: 5F9R^1^. **(b)** The gRNA and target strand of Cas9-ggX20-DNA and Cas9-X20-DNA after aligning the residues of Cas9 between the two structures (Cas9-ggX20-DNA; PDB ID: 5Y36^2^ and Cas9-X20-DNA; PDB ID: 5F9R^1^). With the extensions, the RNA-DNA hybrid at the PAM-distal end has been considerably repositioned, compared with its position with the canonical gRNA. The 21^st^ and 22^nd^ nucleotide, counting from PAM, of the target strand are not base-paired with the non-target strand nucleotides, but are rather flipped towards the gg of the ggX20. **(c)** The relative changes in positions of RNA-DNA hybrid and Cas9-HF1 mutations between Cas9-X20-DNA and Cas9-ggX20-DNA.

## REFERENCES

1. Hille, F. et al. The Biology of CRISPR-Cas: Backward and Forward. Cell 172, 1239–1259 (2018).

2. Knott, G. J. & Doudna, J. A. CRISPR-Cas guides the future of genetic engineering. Science (80-.). 361, 866–869 (2018).

3. Jinek, M. et al. A Programmable Dual-RNA-Guided DNA Endonuclease in Adaptive Bacterial Immunity. Science (80-.). 337, 816–821 (2012).

4. Gasiunas, G., Barrangou, R., Horvath, P. & Siksnys, V. Cas9-crRNA ribonucleoprotein complex mediates specific DNA cleavage for adaptive immunity in bacteria. Proc. Natl. Acad. Sci. 109, E2579–E2586 (2012).

5. Sternberg, S. H., Redding, S., Jinek, M., Greene, E. C. & Doudna, J. A. DNA interrogation by the CRISPR RNA-guided endonuclease Cas9. Nature 507, 62–67 (2014).

6. Szczelkun, M. D. et al. Direct observation of R-loop formation by single RNA-guided Cas9 and Cascade effector complexes. Proc. Natl. Acad. Sci. 111, 9798–9803 (2014).

7. Singh, D., Sternberg, S. H., Fei, J., Doudna, J. A. & Ha, T. Real-time observation of DNA recognition and rejection by the RNA-guided endonuclease Cas9. Nat. Commun. 7, 12778 (2016).

8. Singh, D. & Ha, T. Understanding the Molecular Mechanisms of the CRISPR Toolbox Using Single Molecule Approaches. ACS Chem. Biol. (2018). doi:10.1021/acschembio.7b00905

9. Sternberg, S. H., LaFrance, B., Kaplan, M. & Doudna, J. A. Conformational control of DNA target cleavage by CRISPR–Cas9. Nature 527, 110–113 (2015).

10. Slaymaker, I. M. et al. Rationally engineered Cas9 nucleases with improved specificity. Science 351, 84–8 (2016).

11. Kleinstiver, B. P. et al. High-fidelity CRISPR–Cas9 nucleases with no detectable genome-wide off-target effects. Nature 529, 490–495 (2016).

12. Chen, J. S. et al. Enhanced proofreading governs CRISPR-Cas9 targeting accuracy. Nature 550, 407–410 (2017).

13. Casini, A. et al. A highly specific SpCas9 variant is identified by in vivo screening in yeast. Nat. Biotechnol. 36, 265–271 (2018).

14. Hu, J. H. et al. Evolved Cas9 variants with broad PAM compatibility and high DNA specificity. Nature 556, 57–63 (2018).

15. Lee, J. K. et al. Directed evolution of CRISPR-Cas9 to increase its specificity. Nat. Commun. 9, 3048 (2018).

16. Nishimasu, H. et al. Engineered CRISPR-Cas9 nuclease with expanded targeting space. Science (80-.). eaas9129 (2018). doi:10.1126/science.aas9129

17. Vakulskas, C. A. et al. A high-fidelity Cas9 mutant delivered as a ribonucleoprotein complex enables efficient gene editing in human hematopoietic stem and progenitor cells. Nat. Med. 24, 1216–1224 (2018).

18. Singh, D. et al. Mechanisms of improved specificity of engineered Cas9s revealed by single-molecule FRET analysis. Nat. Struct. Mol. Biol. 25, 347–354 (2018).

19. Kleinstiver, B. P. et al. High-fidelity CRISPR–Cas9 nucleases with no detectable genome-wide off-target effects. Nature 529, 490–495 (2016).

20. Fu, Y., Sander, J. D., Reyon, D., Cascio, V. M. & Joung, J. K. Improving CRISPR-Cas nuclease specificity using truncated guide RNAs. Nat. Biotechnol. 32, 279–284 (2014).

21. Sung, Y. H. et al. Highly efficient gene knockout in mice and zebrafish with RNA-guided endonucleases. Genome Res. 24, 125–31 (2014).

22. Kim, S., Kim, D., Cho, S. W., Kim, J. & Kim, J.-S. Highly efficient RNA-guided genome editing in human cells via delivery of purified Cas9 ribonucleoproteins. Genome Res. 24, 1012–9 (2014).

23. Ha, T. et al. Probing the interaction between two single molecules: fluorescence resonance energy transfer between a single donor and a single acceptor. Proc. Natl. Acad. Sci. U. S. A. 93, 6264–8 (1996).

24. Roy, R., Hohng, S. & Ha, T. A practical guide to single-molecule FRET. Nat. Methods 5, 507–516 (2008).

25. Ivanov, I. E. et al. CAS9 INTERROGATES DNA IN DISCRETE STEPS MODULATED BY MISMATCHES AND SUPERCOILING. in Biophysical Society Meeting (Cell Press, 2019).

26. Oakley, J. L. & Coleman, J. E. Structure of a promoter for T7 RNA polymerase. Proc. Natl. Acad. Sci. U. S. A. 74, 4266–70 (1977).

27. Nishimasu, H. et al. Crystal Structure of Cas9 in Complex with Guide RNA and Target DNA. Cell 156, 935–949 (2014).

28. Huai, C. et al. Structural insights into DNA cleavage activation of CRISPR-Cas9 system. Nat. Commun. 8, 1375 (2017).

29. Dagdas, Y. S., Chen, J. S., Sternberg, S. H., Doudna, J. A. & Yildiz, A. A conformational checkpoint between DNA binding and cleavage by CRISPR-Cas9. Sci. Adv. 3, eaao0027 (2017).

30. Jiang, F. et al. Structures of a CRISPR-Cas9 R-loop complex primed for DNA cleavage. Science 351 867–71 (2016).

31. Singh, D. et al. Mechanisms of improved specificity of engineered Cas9s revealed by single-molecule FRET analysis. Nat. Struct. Mol. Biol. 25, 347–354 (2018).

32. Huai, C. et al. Structural insights into DNA cleavage activation of CRISPR-Cas9 system. Nat. Commun. 8, 1375 (2017).

33. Yin, H. et al. Partial DNA-guided Cas9 enables genome editing with reduced off-target activity. Nat. Chem. Biol. 14, 311–316 (2018).

34. Cromwell, C. R. et al. Incorporation of bridged nucleic acids into CRISPR RNAs improves Cas9 endonuclease specificity. Nat. Commun. 9, 1448 (2018).

35. Rueda, F. O. et al. Mapping the sugar dependency for rational generation of a DNA-RNA hybrid- guided Cas9 endonuclease. Nat. Commun. 8, 1610 (2017).

36. Guschin, D. Y. et al. A Rapid and General Assay for Monitoring Endogenous Gene Modification. in Methods in molecular biology (Clifton, N.J.) 649, 247–256 (2010).

37. Sander, J. D. & Joung, J. K. CRISPR-Cas systems for editing, regulating and targeting genomes. Nat. Biotechnol. 32, 347–355 (2014).

38. Clarke, R. et al. Enhanced Bacterial Immunity and Mammalian Genome Editing via RNA-Polymerase-Mediated Dislodging of Cas9 from Double-Strand DNA Breaks. Mol. Cell 71, 42–55.e8 (2018).

39. Kato-Inui, T., Takahashi, G., Hsu, S. & Miyaoka, Y. Clustered regularly interspaced short palindromic repeats (CRISPR)/CRISPR-associated protein 9 with improved proof-reading enhances homology-directed repair. Nucleic Acids Res. 46, 4677–4688 (2018).

## REFERENCES

1. Jiang, F. et al. Structures of a CRISPR-Cas9 R-loop complex primed for DNA cleavage. Science 351, 867–71 (2016).

2. Huai, C. et al. Structural insights into DNA cleavage activation of CRISPR-Cas9 system. Nat. Commun. 8, 1375 (2017).

