## Supplementary Information for "Single molecule analysis of effects of non-canonical guide RNAs and specificity-enhancing mutations on Cas9-induced DNA unwinding"

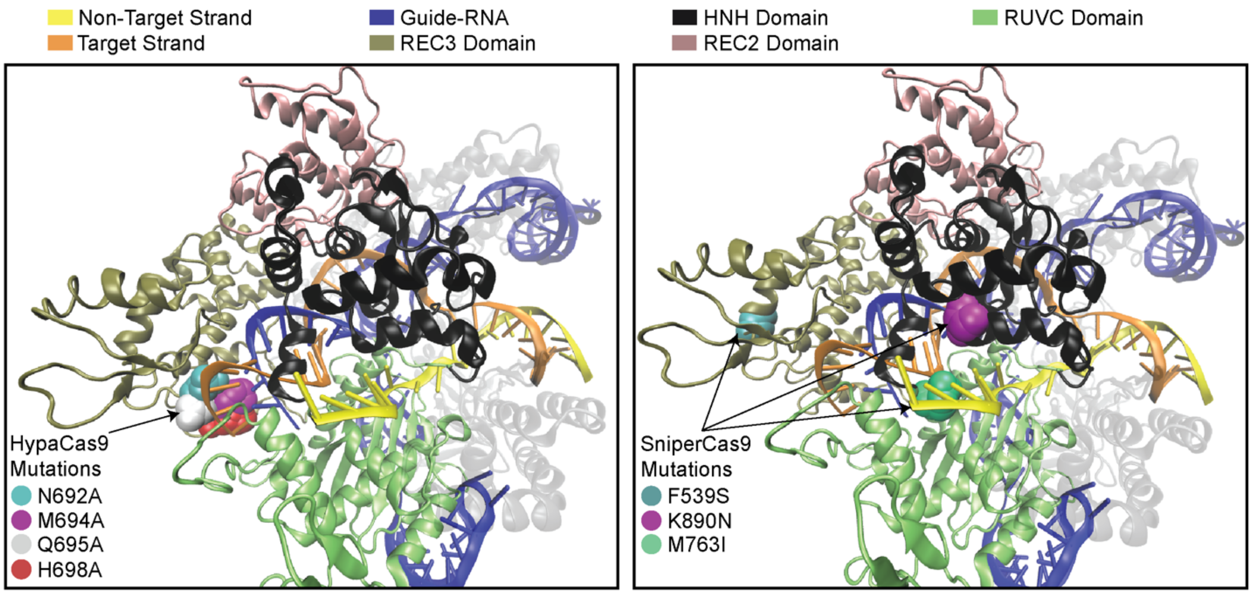


**Supplementary Figure 1. Locations of HypaCas9 and SniperCas9 mutations in the Cas9-RNA-DNA.** (PDB ID: 5F9R[1]).

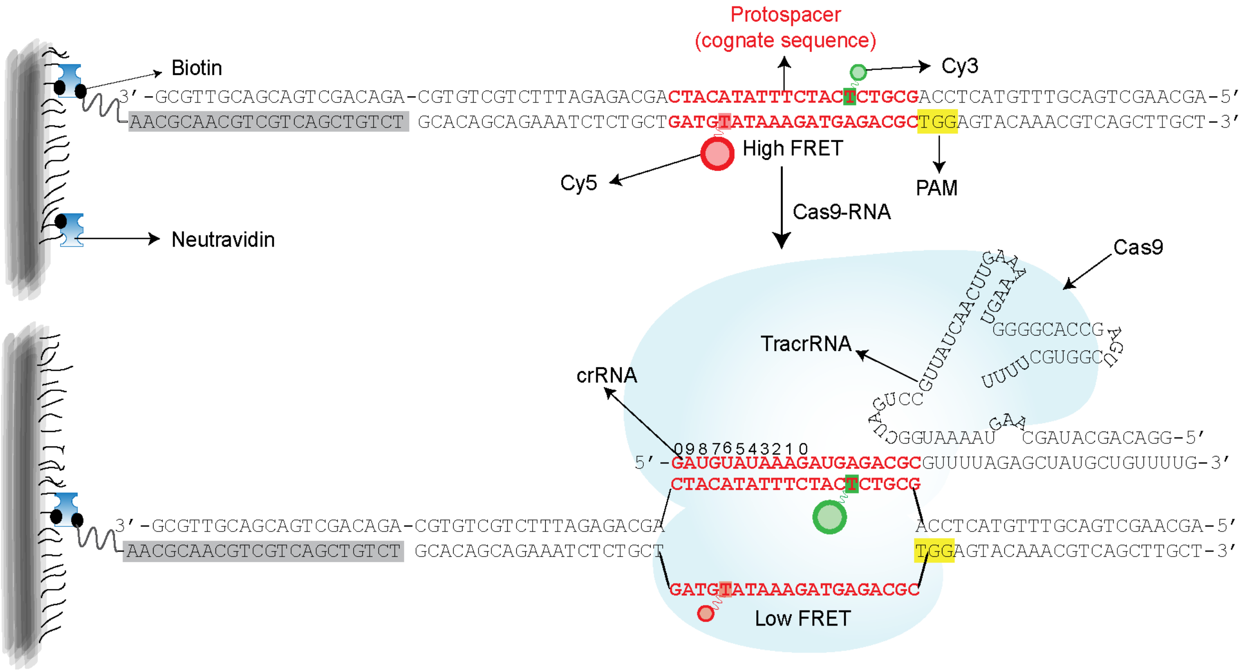


**Supplementary Figure 2. Locations of FRET probes for smFRET DNA unwinding experiments and DNA unwinding by WT Cas9 compared with EngCas9.** Schematic of Cas9-RNA-DNA complex. crRNA hybridized with tracrRNA is the guide-RNA, and the denoted red sequences hybridize with one of the strands (target strand) in the DNA. The strand complementary to target strand in the DNA is the non-target strand which also contains the PAM (5'-NGG-3') as indicated. Highlighted in grey is a 22 nt biotin-labeled strand which is used as an adaptor for surface immobilization of the DNA. The donor and acceptor labels are conjugated to the 6$^{th}$ nucleotide (from PAM) in target strand and 16$^{th}$ in the non-target strand. Cas9-RNA binds the DNA leading to the unwinding of the DNA which results in an increase in distance between FRET labels and thus a decrease in *E*.

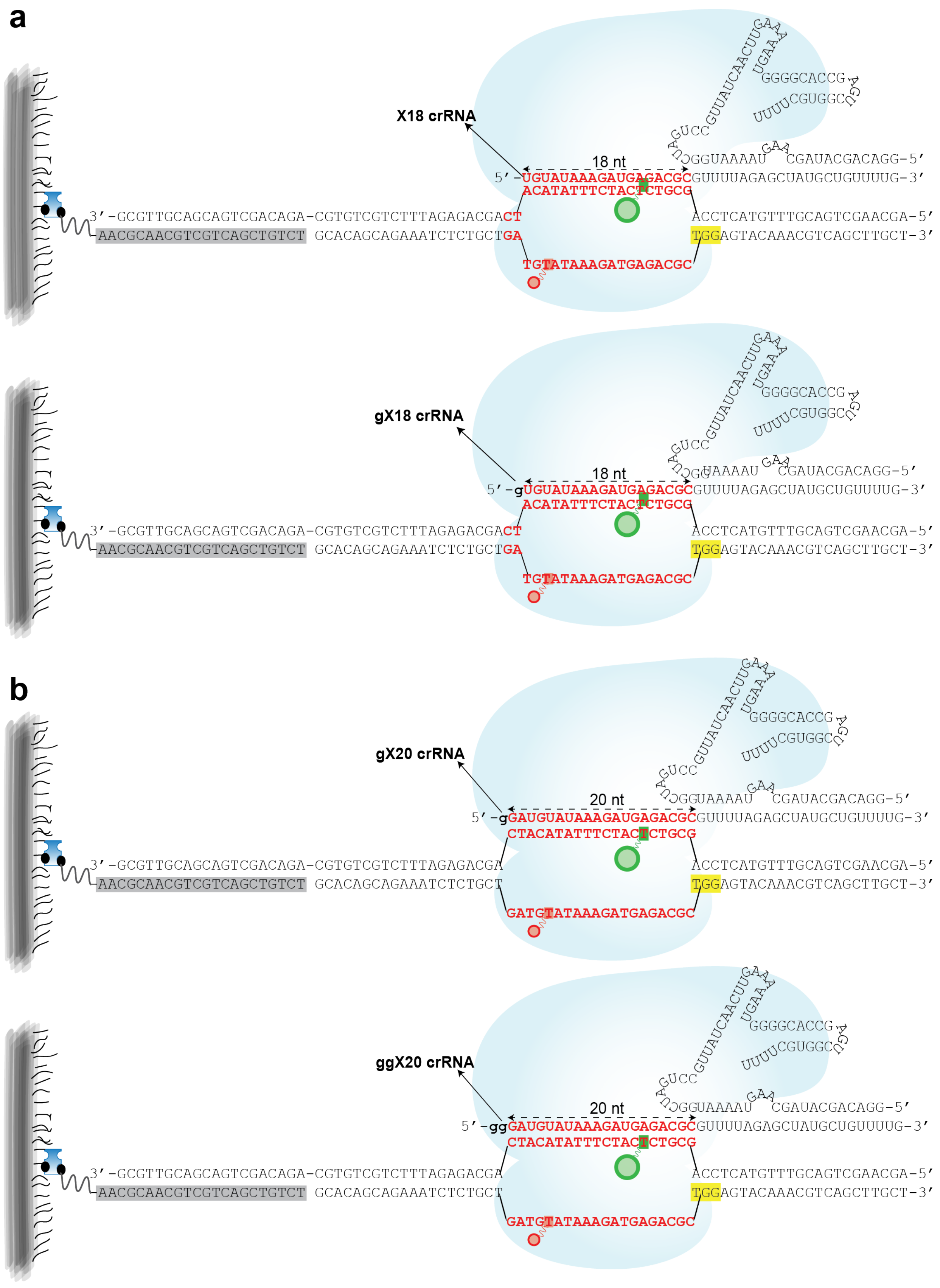

a
X18 crRNA
18 nt
5′-UGUAUAAAGAUGAGACGCGUUUUAGAGCUAUGCUGUUUUG-3′
ACATATTTCTACTCTGCG
3′-GCGTTGCAGCAGTCGACAGA-CGTGTCGTCTTTAGAGACGACT
AACGCAACGTCGTCAGCTGTCT GCACAGCAGAAATCTCTGCTGA
TGTATAAAGATGAGACGC
ACCTCATGTTTGCAGTCGAACGA-5′
TGGAGTACAAACGTCAGCTTGCT-3′
CGAUACGACAGG-5′
gX18 crRNA
18 nt
5′-gUGUAUAAAGAUGAGACGCGUUUUAGAGCUAUGCUGUUUUG-3′
b
gX20 crRNA
20 nt
5′-gGAUGUAUAAAGAUGAGACGCGUUUUAGAGCUAUGCUGUUUUG-3′
CTACATATTTCTACTCTGCG
GATGTATAAAGATGAGACGC
ggX20 crRNA
20 nt
5′-ggGAUGUAUAAAGAUGAGACGCGUUUUAGAGCUAUGCUGUUUUG-3′

**Supplementary Figure 3. Schematic of the smFRET unwinding assay involving non-canonical gRNAs.** (a) Extended gRNAs. (b) Truncated gRNAs.

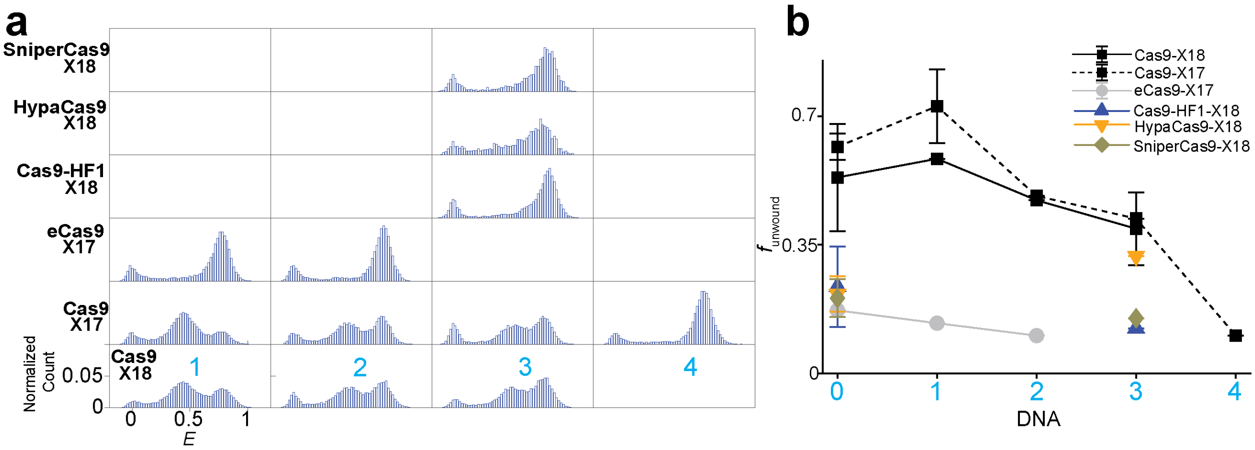


**Supplementary Figure 4. Unwinding by truncated gRNA.**

**(a)** *E* histograms of unwinding by truncated gRNA. Each row corresponds to a particular labeled Cas9 and gRNA combination. Each column corresponds to a particular DNA target. **(b)** $f_{\text{unwound}}$ vs. $n_{\text{PD}}$ for different Cas9s with truncated gRNAs.

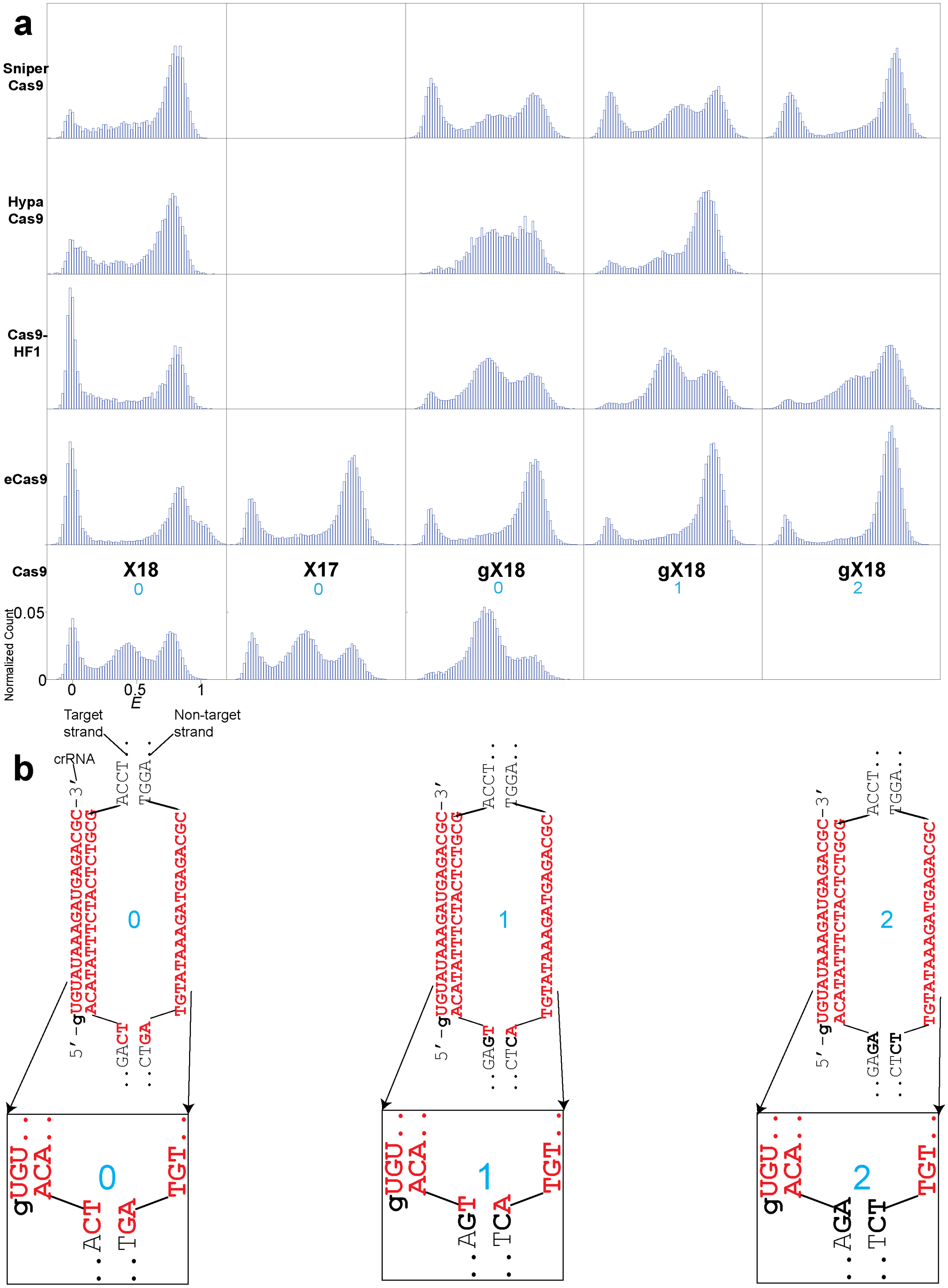
a
Sniper Cas9
Hypa Cas9
Cas9-HF1
eCas9
Cas9
X18
0
X17
0
gX18
0
gX18
1
gX18
2
Normalized Count
0.05
0
0
0.5
1
E
b
Target strand
Non-target strand
crRNA
5′-gUGUAUAAAGAUGAGACGC-3′
ACATATTTCTACTCTGCG
TGTATAAAGATGAGACGC
0
1
2

**Supplementary Figure 5. Unwinding by truncated gRNA.**

**(a)** *E* histograms of unwinding by truncated gRNA. Each row corresponds to a particular labeled Cas9. Each column corresponds to a particular gRNA and DNA target. **(b)** Sequences of DNA and crRNA (of truncated gRNAs). X18/gX18 guide RNA forms the same Watson-Crick base pairing with $n_{PD}$=0, $n_{PD}$=1, and $n_{PD}$=2 DNA targets.

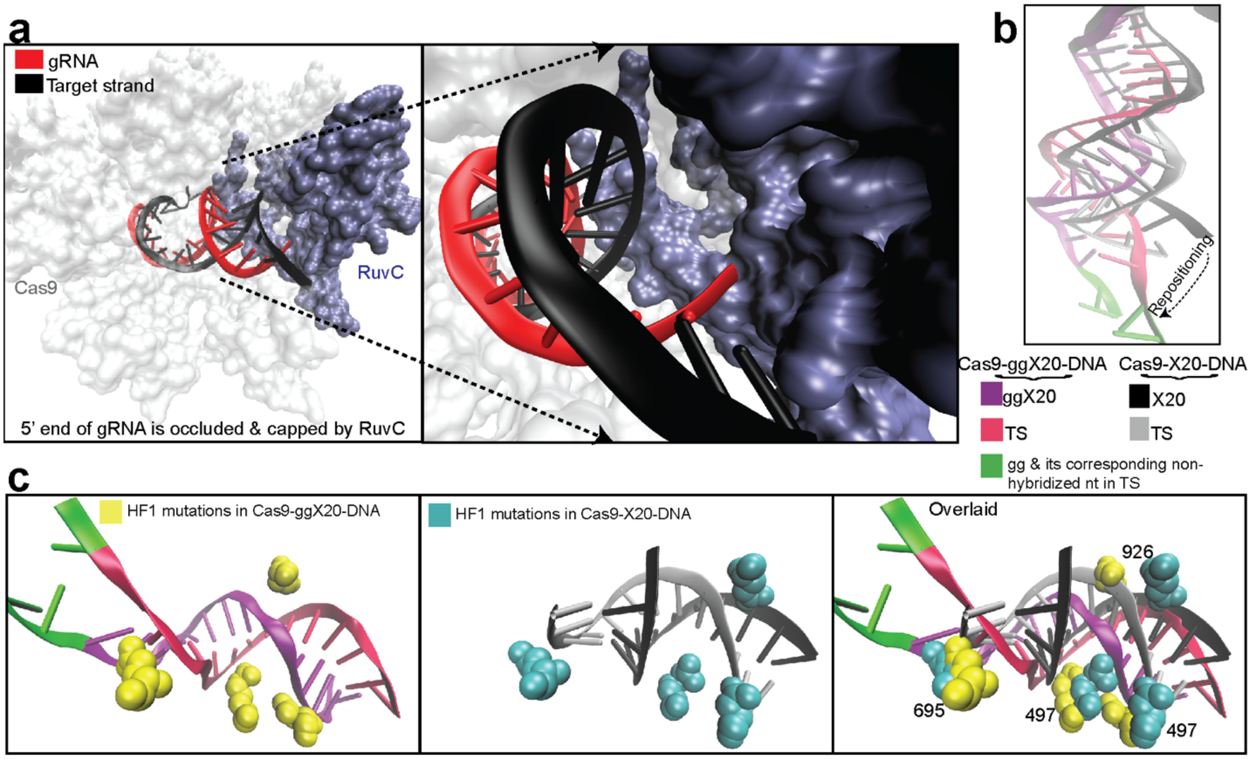


**Supplementary Figure 6. PAM distal end of Cas9-RNA-DNA complex.**

**(a)** 5' end of the gRNA is occluded and capped by the RuvC nuclease domain in Cas9-X20-DNA. PDB ID: 5F9R[1]. **(b)** The gRNA and target strand of Cas9-ggX20-DNA and Cas9-X20-DNA after aligning the residues of Cas9 between the two structures (Cas9-ggX20-DNA; PDB ID: 5Y36[2] and Cas9-X20-DNA; PDB ID: 5F9R[1] ). With the extensions, the RNA-DNA hybrid at the PAM-distal end has been considerably repositioned, compared with its position with the canonical gRNA. The 21$^{st}$ and 22$^{nd}$ nucleotide, counting from PAM, of the target strand are not base-paired with the non-target strand nucleotides, but are rather flipped towards the gg of the ggX20. **(c)** The relative changes in positions of RNA-DNA hybrid and Cas9-HF1 mutations between Cas9-X20-DNA and Cas9-ggX20-DNA.

**REFERENCES**


1. Jiang, F. *et al.* Structures of a CRISPR-Cas9 R-loop complex primed for DNA cleavage. *Science* **351,** 867–71 (2016).
2. Huai, C. *et al.* Structural insights into DNA cleavage activation of CRISPR-Cas9 system. *Nat. Commun.* **8,** 1375 (2017).
